# Intra- and inter-specific variations of gene expression levels in yeast are largely neutral

**DOI:** 10.1101/089995

**Authors:** Jian-Rong Yang, Calum Maclean, Chungoo Park, Huabin Zhao, Jianzhi Zhang

**Affiliations:** Department of Ecology and Evolutionary Biology, University of Michigan, Ann Arbor, MI 48109, USA

**Keywords:** Evolution, transcriptome, genetic drift, adaptation, *Saccharomyces*

## Abstract

It is commonly, although not universally, accepted that most intra- and inter-specific genome sequence variations are more or less neutral, whereas a large fraction of organism-level phenotypic variations are adaptive. Gene expression levels are molecular phenotypes that bridge the gap between genotypes and corresponding organism-level phenotypes. Yet, it is unknown whether natural variations in gene expression levels are mostly neutral or adaptive. Here we address this fundamental question by genome-wide profiling and comparison of gene expression levels in nine yeast strains belonging to three closely related Saccharomyces species and originating from five different ecological environments. We find that the transcriptome-based clustering of the nine strains approximates the genome sequence-based phylogeny irrespective of their ecological environments. Remarkably, only ∼0.5% of genes exhibit similar expression levels among strains from a common ecological environment, no greater than that among strains with comparable phylogenetic relationships but different environments. These and other observations strongly suggest that most intra- and inter-specific variations in yeast gene expression levels result from the accumulation of random mutations rather than environmental adaptations. This finding has profound implications for understanding the driving force of gene expression evolution, genetic basis of phenotypic adaptation, and general role of stochasticity in evolution.

## SIGNIFICANCE

It is believed that most genomic variations within and between species are more or less neutral, while a large fraction of organism-level phenotypic variations are adaptive. Gene expression bridges the gap between genome sequences and organismal phenotypes, but it is unclear whether variations in gene expression levels are mostly neutral or adaptive. By quantifying the genome-wide gene expression levels in nine yeast strains whose phylogenetic relationships mismatch the relationships of their ecological environments, we find that transcriptome variations can be explained by the accumulation of random mutations but not environmental adaptations. This finding has important implications for understanding the driving force of gene expression evolution, genetic basis of phenotypic adaptation, and general role of stochasticity in evolution.

## INTRODUCTION

Evolutionary biology began from studies of organism-level phenotypes such as morphological, physiological, and behavioral traits. Darwin proposed that these variations, be they intra- or inter-specific, can primarily be explained by the adaptation of organisms to their respective environments (1), a view largely shared by modern biologists (2-4) (but see (5)). However, at the genotype level, molecular evolutionists generally agree that most intra- and inter-specific variations in DNA sequences are more or less neutral (6-8). This contrast between genotypic and phenotypic evolution is not logically inconsistent, because a genotypic change need not result in a phenotypic change or one with an appreciable fitness effect, even though a stably inherited phenotypic difference always requires a genotypic change.

If a genotypic change has a potential phenotypic effect, the realization of this potential usually requires gene expression, regardless of whether the genotypic change occurs in a coding region or a non-coding regulatory region. In other words, gene expression is usually the necessary bridge between genotypes and their corresponding organismal phenotypes. In this context, one ponders whether gene expression level, a molecular phenotype, is more like (organismal) phenotypes or (molecular) genotypes in its evolutionary pattern and mechanism. Specifically, we ask whether variations in gene expression levels within and between species result largely from neutral or adaptive evolution. Unfortunately, there is no unequivocal theoretical answer to this question, because an expression-level change may or may not appreciably impact the organismal phenotype and fitness. The available empirical data in the literature do not provide a clear answer either. For instance, although many presumably adaptive morphological variations in nature have been found to be caused by gene expression changes (9,10), this fact at most suggests that a large fraction of morphological adaptations are due to expression changes, but not that a large fraction of expression differences are adaptive. A neutral model of transcriptome evolution was previously proposed, on the basis of, among other things, an approximately constant rate of transcriptome evolution (11). This observation could, however, be an artifact of using a human microarray to measure gene expression levels in other primates (12). Furthermore, due to the lack of specific predictions of the adaptive hypothesis in the primate study, adaptation and neutrality could not be unambiguously distinguished.

To answer whether variations in gene expression levels within and between species are largely neutral or adaptive, we compared the transcriptomes of nine yeast strains belonging to three closely related species isolated from five different ecological environments. Importantly, we selected strains such that their genomic phylogenetic relationships mismatch their relationships of environmental origins. For instance, some strains are phylogenetically relatively distant from one another but have similar ecological environments, whereas others are phylogenetically relatively close to one another but live in ecologically distinct environments. If gene expression variations among these strains result from the accumulation of neutral mutations, the relationships of their transcriptomes should mimic the genome-based phylogenetic tree. On the contrary, if the expression variations among these strains are largely shaped by adaptations to their respective environments, their transcriptomes should cluster according to their ecological environments. Thus, our design allows a distinction between the neutral and adaptive hypotheses. We quantified the yeast transcriptomes by RNA sequencing (RNA-seq) and discovered that these transcriptomes cluster more or less according to the genome-based phylogeny rather than their ecological environments. Only a tiny fraction of yeast genes exhibit similar expression levels among strains with a common ecological environment, no greater than that among strains with comparable phylogenetic relationships but different environments. Our findings strongly suggest that the vast majority of yeast gene expression variations result from neutral rather than adaptive evolution.

## RESULTS

### The transcriptome tree approximates the genome tree

To study the driving force of gene expression evolution, we chose nine yeast strains belonging to three closely related species (sister species *Saccharomyces cerevisiae* and *S. paradoxus*, plus their outgroup *S. mikatae*) and originating from five different ecological environments (**Fig. 1A**; see also **Table S1**). The natural habitat of yeast is thought to be the sap and bark of oak trees and adjacent soils (13). Five of our nine strains were collected from this ecological environment and are referred to as wild strains. They include two *S. cerevisiae* strains, two *S. paradoxus* strains, and one *S. mikatae* strain. Four additional *S. cerevisiae* strains were respectively isolated from four other environments and are referred to as non-wild strains.

**Figure 1.**
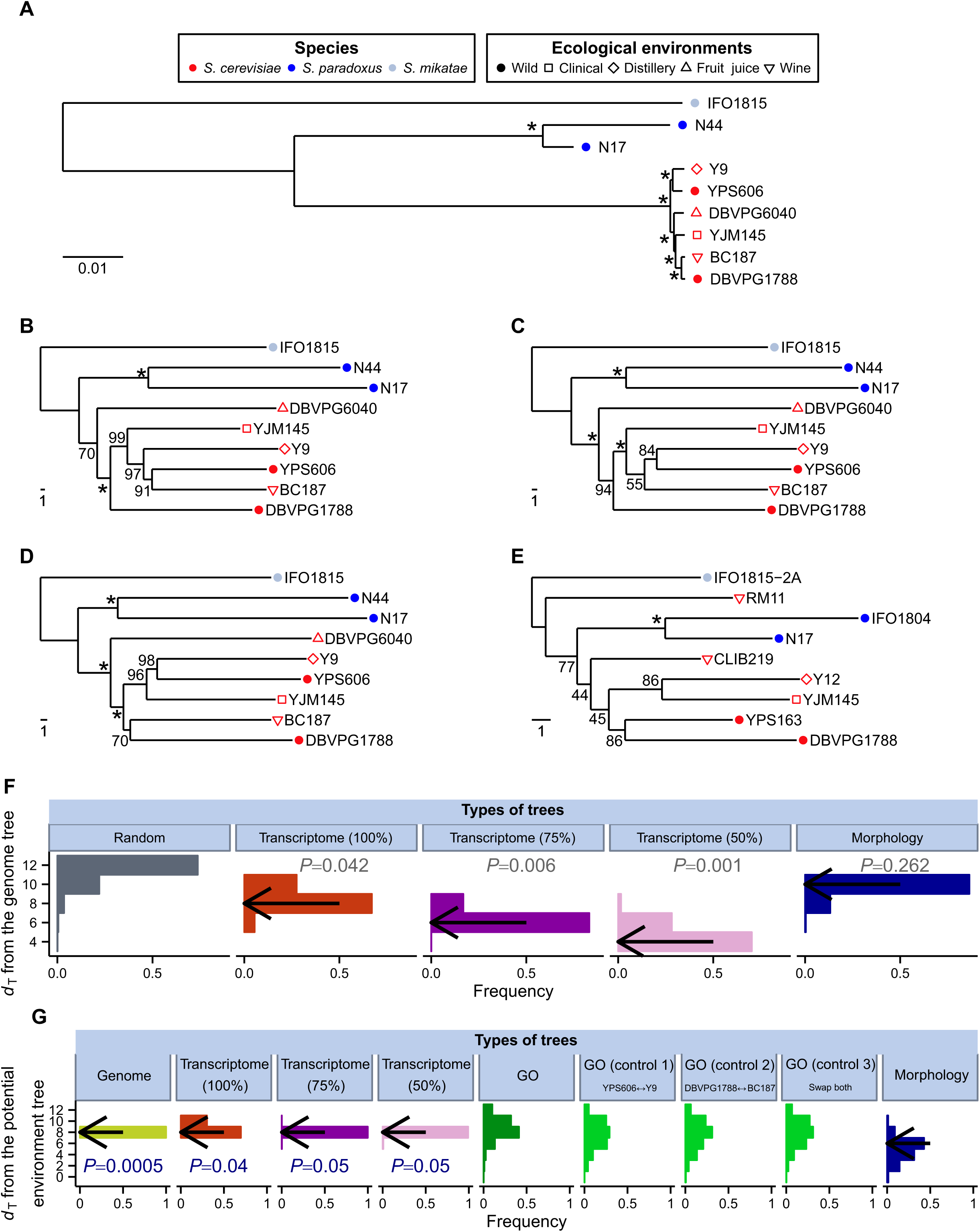
Phylogenetic trees of the nine *Saccharomyces* yeast strains constructed using genome sequence, gene expression, and morphology data, respectively. The three species are indicated by different colors, while the ecological environments where the strains were isolated are shown by different symbols. (A) The genome tree of the nine strains based on the alignment of the coding sequences of 4325 genes. Bootstrap percentages estimated from 1000 replications are shown on interior branches. Asterisks indicate > 99.5% bootstrap support. The scale bar shows 0.01 nucleotide substitutions per site. (B-D) The transcriptome tree of the nine strains based on standardized Euclidian distances in gene expression levels of all 4325 genes (B), the 75% most highly expressed genes (C), and the 50% most highly expressed genes (D). Bootstrap percentages estimated from 10,000 replications are shown on interior branches. Asterisks indicate > 99.5% bootstrap support. The scale bar shows 1 unit of the standardized Euclidian distance. (E) The morphology tree of nine strains based on standardized Euclidian distances in 219 morphological traits. Strains IFO1804, RM11, CLIB219, Y12, and YPS163 are used as proxies of N44, BC187, DBVPG6040, Y9, and YPS606, respectively. Bootstrap percentages estimated from 10,000 replications are shown on interior branches. Asterisks indicate > 99.5% bootstrap support. The scale bar shows 1 unit of the standardized Euclidian distance. (F) Frequency distributions of topological distances (*d*_T_) between the genome tree and random tree topologies (grey), bootstrapped transcriptome trees with all genes (brown), bootstrapped transcriptome trees with the 75% most highly expressed genes (dark purple), bootstrapped transcriptome trees with the 50% most highly expressed genes (light purple), and bootstrapped morphology trees (blue), respectively. Each distribution is based on 10,000 random trees or bootstrapped trees. Arrows indicate the observed *d*_T_ between the genome tree and various other trees based on the original (rather than bootstrapped) data. *P*-value shows the probability with which the *d*_T_ between the genome tree and a random tree topology is smaller than the observed *d*_T_ between the genome tree and the tree being compared. (G) Frequency distributions of topological distances (*d*_T_) between the potential environment tree and bootstrapped genome trees (yellow), bootstrapped transcriptome trees with all genes (brown), bootstrapped transcriptome trees with the 75% most highly expressed genes (dark purple), bootstrapped transcriptome trees with the 50% most highly expressed genes (light purple), gene expression trees based on 533 individual GO categories (dark green), and bootstrapped morphology trees (blue), respectively, as well as frequency distributions of *d*_T_ between three control environment trees and 533 GO-based gene expression trees, respectively (light green). The *d*_T_ between a tree and the potential environment tree is defined by the minimal topological distance between the tree and any tree containing a monophyly of the five wild strains. In the three control environment trees, one or both *S. cerevisiae* wild strains in the aforementioned monophyly are swapped with their sister strains in the genome tree. Each distribution except for the bootstrapped genome trees (1000 replications) and GO-based trees (533 GO categories) is derived from 10,000 bootstrapped trees. Arrows indicate the observed *d*_T_ between the potential environment tree and various other trees based on the original (rather than bootstrapped) data.

On the basis of the available genome sequences of these nine strains, we built a multiple sequence alignment of the coding sequences of each of 4325 genes that have fully sequenced and reliable one-to-one orthologs across the nine strains (see Materials and Methods). We concatenated the aligned sequences after removing all gaps and used them to reconstruct a neighbor-joining (NJ) tree (**Fig. 1A**), which will be referred to as the genome tree. This tree is highly resolved, with all nodes having > 99.5% bootstrap support. As expected, the nine strains are clustered in the genome tree by species identity rather than ecological environment. Furthermore, within the *S. cerevisiae* clade, the two wild strains each have a non-wild sister strain. If gene expression variations among the nine strains are in a large part due to the accumulation of neutral mutations, the transcriptomes of these strains should cluster in a tree similar to the genome tree. If gene expression variations among the strains are largely caused by environmental adaptations, the five wild strains should cluster in the transcriptome tree, contrasting the genome tree.

To distinguish between the above two hypotheses, we quantified the genome-wide gene expression levels of the nine strains in the same growth medium and growth phase such that the revealed expression variations reflect genetic differences rather than phenotypic plasticity. The synthetic medium used mimics oak exudate (14), rendering potential expression adaptations of the wild strains readily detectable. We used diploid strains, as yeast is naturally homothallic (13, 15, 16) and therefore usually diploid because gametes can switch mating type after dividing and mate with their daughter cells. RNAs were extracted from one exponentially dividing culture of each of the nine strains and a replicate culture of the *S. cerevisiae* wild strain YPS606 during exponential growth, and the standard Illumina-based RNA-seq was performed (see Materials and Methods). A total of ∼744 million 52-nt single-end reads were obtained (**Table S1**); these reads were mapped to their respective genomes and used to estimate gene expression levels (see Materials and Methods). Our experiments generated highly reproducible results, because gene expression levels quantified in the two biological replicates of strain YPS606 have a Pearson correlation coefficient of 0.984 (*P* < 10^−300^; **Fig. S1**). These two replicates were subsequently pooled in our analysis unless otherwise noted.

Our transcriptome analysis focused on the same 4325 genes used to reconstruct the genome tree, each of which contains at least one read in at least one of the nine strains. Using the transcriptome data, we calculated the standardized Euclidian distance between each pair of the nine strains (see Materials and Methods) and then built an NJ tree, referred to as the transcriptome tree (**Fig. 1B**). The overall topology of the transcriptome tree is similar but not identical to that of the genome tree. Importantly, we found the nine strains to cluster in the transcriptome tree by species identity rather than ecological environment. Although the transcriptome tree shows a different topology from that of the genome tree for the six *S. cerevisiae* strains, the two *S. cerevisiae* wild strains are not clustered in the transcriptome tree (**Fig. 1B**). Thus, the intra-specific topological differences between the transcriptome tree and genome tree do not appear to reflect environmental adaptations in gene expression evolution. When we separately analyzed the RNA-seq data from the two biological replicates of the YPS606 strains, the two replicates cluster in the transcriptome tree, as expected (**Fig. S2**).

Despite the high sequencing depths of our RNA-seq experiments, expression level estimates of lowly expressed genes are still less reliable than those of other genes. Because the standardized Euclidian distance gives equal weights to all genes, including lowly expressed genes in the analysis increases the sampling error of the transcriptome tree. We thus repeated the above analysis using only the 75% (**Fig. 1C**) or 50% (**Fig. 1D**) most highly expressed genes after ranking all genes by their mean expression levels across all strains. Interestingly, using these relatively highly expressed genes renders the transcriptome trees even more similar to the genome tree. Specifically, they recovered the genome-tree-based relationships among some *S. cerevisiae* strains. This is not unexpected, given that (i) the differences between the transcriptome tree made using all genes and the genome tree seem random and (ii) excluding lowly expressed genes should increase the signal to noise ratio for making the transcriptome tree.

To quantify the topological differences between the transcriptome tree made using all genes and the genome tree, we measured their topological distance (*d*_T_). The *d*_T_ between two unrooted trees is twice the number of interior branches at which taxon partition is different between the two trees compared (see Materials and Methods). We found that *d*_T_ = 8 between the genome tree and transcriptome tree. In comparison, we generated 10,000 random tree topologies among the nine strains (see Materials and Methods) and calculated their *d*_T_ from the genome tree. We found that only 4.2% of these random topologies have a *d*_T_ ≤ 8 (**Fig. 1F**), suggesting that the small *d*_T_ between the transcriptome tree and genome tree is unlikely to have been caused by chance. The *d*_T_ values from the genome tree reduce to 6 and 4 for the transcriptome trees built using the 75% and 50% most highly expressed genes, respectively, and these *d*_T_ values are again significantly smaller than expected by chance (*P* = 0.006 and 0.001, respectively; (**Fig. 1F**).

The exact topology of the environment tree that describes the relative similarities among the ecological environments of the nine strains is unknown, except that the tree should contain a monophyly of the five wild strains. We thus defined an environment tree set by all trees that satisfy the above condition. The *d*_T_ between the genome tree and the potential environment tree was defined by the smallest topological distance between the genome tree and any tree in the environment tree set. We similarly defined the *d*_T_ between the transcriptome tree and the potential environment tree. We found no difference between these two *d*_T_ values (**Fig. 1G**), indicating that the transcriptome tree is not closer than the genome tree to the potential environment tree. The same is true when only the 75% or 50% most strongly expressed genes are considered (**Fig. 1G**).

While the above analyses found no evidence for the adaptive hypothesis of transcriptome evolution, it is important to confirm that our experimental design is able to detect adaptations if they exist. In this context, it is worth mentioning a yeast phenome study, which measured three growth characteristics in 200 different conditions for a number of strains and then clustered the strains by similarity in their growth characteristics (17). Four of the five wild strains studied here (except DBVPG1788) form a monophyly in the growth traits-based tree (17), suggesting that phylogenetic clustering is able to detect at least some potential adaptations. As a further verification, we analyzed 219 morphological traits previously measured from fluorescent microscopic images of triple-stained yeast cells (18, 19). Using this dataset and controlling for mutational size, we recently discovered that morphological traits that are more important to organismal fitness tend to differ more between strains, strongly suggesting that the intra- and inter-specific morphological variations in this dataset have been shaped by adaptive evolution to a large extent (19). We thus subjected the yeast morphological data to the same phylogenetic analysis used for the transcriptome data. However, five of the nine strains with transcriptome data do not have morphological data. We thus chose five other strains that have morphological data as their proxies. Each of the proxies is ecologically equivalent (with one exception) and, based on the established genome trees (20, 21), phylogenetically close to the strain being replaced. The exception is the *S. cerevisiae* fruit juice strain DBVPG6040. Because this strain has no phylogenetically close and ecologically equivalent strain in the set of strains with morphological data, we chose a phylogenetically close wine strain (CLIB219) as its proxy. We then calculated standardized Euclidian distances between pairs of the nine strains using all 219 morphological traits and built an NJ tree (**Fig. 1E**). Compared with the transcriptome tree, the morphology tree is more different in overall topology from the genome tree. For instance, morphological clustering of the strains is no longer strictly by species identity, because the six *S. cerevisiae* strains do not form a monophyletic clade. Furthermore, in contrast to the genome tree and transcriptome tree, the morphology tree unites the two wild *S. cerevisiae* strains in exclusion of all other strains. We found the *d*_T_ between the morphology tree and genome tree to be 10, not significantly smaller than that between a random tree and the genome tree (*P* = 0.262; **Fig. 1F**). This is not caused by the existence of two wine strains in the morphological data, because these two strains are not clustered in the tree. The *d*_T_ between the morphology tree and potential environment tree is smaller than that between the genome tree and potential environment tree (**Fig. 1G**). To examine if this difference is statistically significant, we generated 10,000 morphology trees and 10,000 genome trees by bootstrapping the 219 morphological traits and the 4325 genes, respectively. The mean *d*_T_ to the potential environment tree is significantly smaller for the set of morphology bootstrap trees than the set of genome bootstrap trees (*P* = 0.0005, *Z-*test; **Fig. 1G**). These observations confirm that our experimental design is able to detect potential signals of environmental adaptation.

We similarly used the bootstrap method with 10,000 replications to examine if the *d*_T_ between the transcriptome tree and genome tree is significantly smaller than that between the morphology tree and genome tree. While this difference in *d*_T_ is not statistically significant (*P* = 0.16; **Fig. 1F**), the difference becomes significant when only the 75% or 50% most highly expressed genes are used in the transcriptome analysis (*P* = 6×10^−4^ and 2×10^−5^, respectively; **Fig. 1F**). By the same approach, we found that the *d*_T_ between any of the three transcriptome trees and the potential environment tree is significantly larger than that between the morphology tree and potential environment tree (*P* = 0.04, 0.05, and 0.05, respectively; **Fig. 1G**). These results further support the hypothesis of neutral rather than adaptive evolution of yeast transcriptomes.

### Principal component analysis supports the phylogenetic results

In addition to the phylogenetic analysis, we performed a principal component analysis (PCA) of the genome sequence, transcriptome, and morphology data, respectively (see Materials and Methods). For the genome sequence data, the nine strains are clearly separated according to species identity rather than ecological environment in the plane of the first two principal components (**Fig. 2A**). Although to a lesser extent, the same can be said for the transcriptome data (**Fig. 2B**; **Figs. S3A** and **B**). By contrast, there is no clear grouping of strains by species identity when the morphological data are analyzed (**Fig. 2C**). For instance, the distances between some pairs of inter-specific strains on the PCA plot are smaller than the distances between some pairs of intra-specific strains. Of special interest is that the distance between the two *S. cerevisiae* wild strains is much smaller than that between each of them and their respective sister strains or their proxies. The contrast between the morphology PCA plot and transcriptome PCA plot is not due to the much larger number of genes/traits in the transcriptome data, compared with that in the morphology data. This is because, even when only 219 randomly picked genes are used, the transcriptome PCA plot still displays a much stronger grouping of strains by species than does the morphology PCA plot (**Fig. S3C**). In summary, results from the PCA and phylogenetic analysis are consistent, both suggesting that the transcriptome variation among the nine strains is largely neutral.

**Figure 2.**
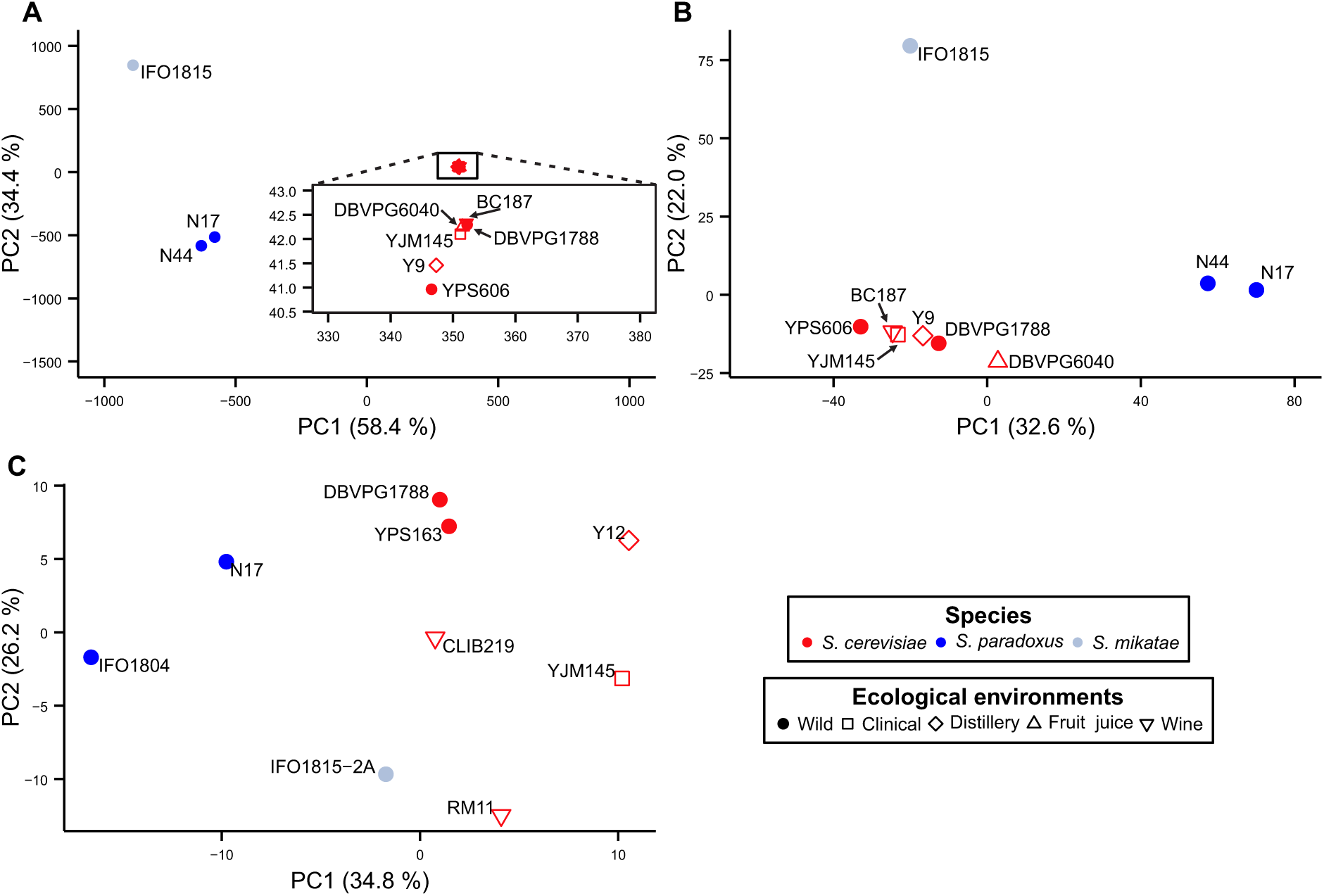
Principal component analysis of the (A) genome sequences, (B) gene expression levels, and (C) morphological data of the nine yeast strains. The three species are indicated by different colors, while the ecological environments where the strains were isolated are shown by different symbols. Percentage variance explained by a principal component is indicated in the parentheses. In panel A, the inset shows an enlarged view of the boxed area.

### Expression variations of functional groups of genes are consistent with neutral evolution

The above finding that the evolution of the transcriptome as a whole resembles genome sequence evolution suggests that the expression variations of most genes are likely neutral. However, this finding does not rule out the possibility that the expression variations of a minority of genes are caused by environmental adaptations. Specifically, these genes may be enriched in certain functional groups. To investigate this possibility, we downloaded the Gene Ontology (GO)-based gene functional annotations in GOslim (22). For each molecular function, biological process, or cellular component GO category, we constructed an NJ tree using the expression levels of all genes belonging to the GO category as was done for all genes in Fig. 1B. Of 533 GO categories examined, only one exhibits a monophyly of the set of all five wild strains. To examine if this observation is explainable by chance alone, we constructed three five-strain control sets by swapping one or both of the two wild *S. cerevisiae* strains with their respective non-wild sister strains shown in the genome tree. We found 3 (swapping DBVPG1788 with BC187), 1 (swapping YPS606 with Y9), and 1 (swapping both) GO categories for which the three five-strain control sets form a monophyly in the expression tree, respectively. Because the observed number (1) for the all-wild strain set is not larger than those (1-3) for the three control sets, we conclude that genes with adaptive expression variations, if they exist, are not significantly enriched in any GO category, which could happen if the number of such genes is small and/or these genes are more or less evenly distributed among GO categories. We further generated the distribution of *d*_T_ between the expression tree of a GO category and the potential environment tree using all 533 GO categories (**Fig. 1G**). It is clear that there is no enrichment of GOs with small *d*_T_ in this distribution, compared with the corresponding distributions when one or both of the two wild *S. cerevisiae* strains are swapped with their respective genomic sister strains in expression trees (**Fig. 1G**).

### Expression variations of individual genes are consistent with neutral evolution

The lack of significant GO enrichment of genes with potential adaptive expression variations prompted us to examine individual genes. For each gene, we calculated the standardized Euclidian distance between each pair of the nine strains, which is simply the absolute value of their standardized expression level difference for the gene (see Materials and Methods). We then used these distances to construct an expression tree by the NJ method. We were interested in expression trees in which the five wild strains form a monophyly, because such trees potentially result from adaptive expression evolution. Only 22 genes met this requirement. A careful examination of their expression variations, however, led to three unexpected observations. First, for each of these 22 genes, all five wild strains showed either higher or lower expressions than all four non-wild strains (**Fig. 3A**). This is unexpected, because the five wild strains could also form a monophyly if they all have similar expression levels that are intermediate to those of non-wild strains, with some non-wild strains having higher expressions and others lower expressions. Second, if the expression variation of a gene is primarily due to environmental adaptations, given the first observation, each of the five wild strains should have the same probability to be the strain with the most dissimilar expression from those of the four non-wild strains. Surprisingly, for only one of the 22 genes is a wild *S. cerevisiae* strain’s expression most dissimilar from those of the non-wild strains, significantly below the expectation of 22×(2/5)=8.8 (*P* = 2×10^−4^, binomial test). Because all four non-wild strains belong to *S. cerevisiae*, the first two observations together strongly suggest that, even for these 22 genes, the vast majority still show a clear signal of expression clustering by species (**Fig. 3A**), which is consistent with neutral evolution. Third, the *c*oefficient of *v*ariation (CV) of gene expression among the six *S. cerevisiae* strains is smaller than the CV among the five wild strains for 15 of the 22 genes. Because the six *S. cerevisiae* strains are from five different environments whereas the five wild strains are all from one environment, the third observation is unexplainable by the environmental adaptation hypothesis but is consistent with the neutral hypothesis. Thus, the expression variations of even these 22 genes are unlikely to have been primarily caused by environmental adaptations.

**Figure 3.**
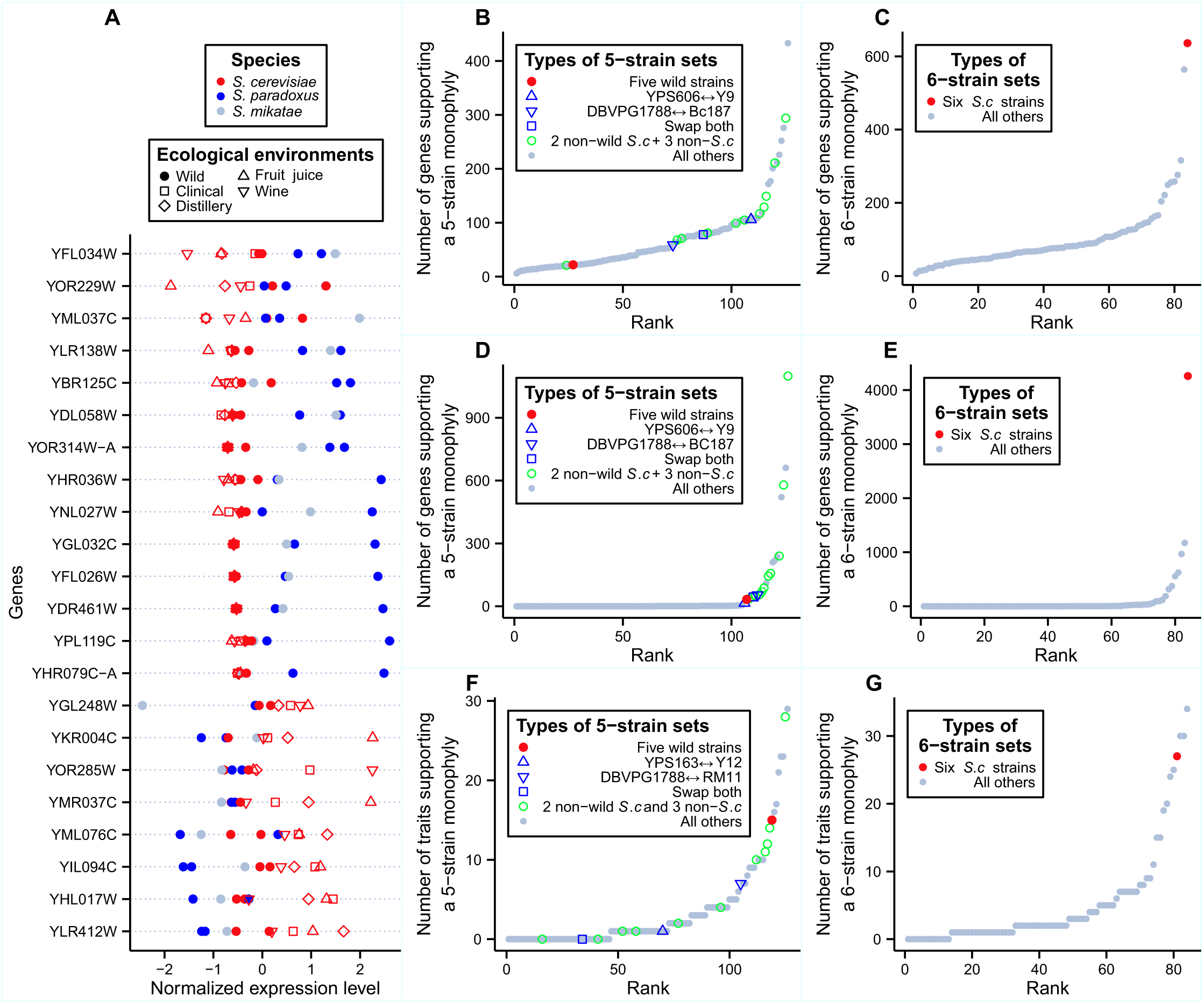
Little evidence for environmental adaptation from the expression levels of individual genes. (A) Twenty-two genes whose expression levels support a monophyly of the five wild strains. Expression levels of the nine strains for each gene have been scaled to a standard normal distribution for comparison. The three species are indicated by different colors, while the ecological environments where the strains were isolated are shown by different symbols. (B) Number of genes whose expression tree supports the monophyly of each of the 126 possible five-strain sets. The red dot shows the all-wild set, the three blue symbols show the three control sets in which one or both of the two wild *S. cerevisiae* strains in the all-wild set are replaced with their sister non-wild strains, the green circles show all other five-strain sets that include the three non-*S. cerevisiae* strains, and the grey dots show all other five-strain sets. (C) Number of genes whose expression tree supports the monophyly of each of the 84 possible six-strain sets. The red dot shows the all-*S. cerevisiae* set, while the grey dots show all other six-strain sets. (D) Same as B except that coding sequence data instead of gene expression data are used. (E) Same as C except that coding sequence data instead of gene expression data are used. (F) Same as B except that morphological data instead of gene expression data are used. (G) Same as C except that morphological data instead of gene expression data are used.

To investigate the possibility that the clustering of the five wild strains by the expression levels for these 22 genes is by chance, we enumerated all 126 ways that five strains can be chosen from the nine strains. For each set of five strains chosen, we calculated the number of genes whose expression trees show a monophyly of these five strains, and referred to this number as the number of monophyly genes (*N*_MG_) for the five-strain set. We ranked *N*_MG_ for the 126 five-strain sets and found that 100 of the 126 sets have *N*_MG_ ≥ 22 (**Fig. 3B**). Furthermore, the all-wild set (red dot in **Fig. 3B**) has the second smallest *N*_MG_ among the 15 five-strain sets that are composed of any two *S. cerevisiae* strains and the three non-*S. cerevisiae* strains (green and blue symbols in **Fig. 3B**), suggesting that non-wild *S. cerevisiae* strains cluster more often with (the wild strains of) the other two species than the wild *S. cerevisiae* strains at the level of gene expression. In addition, we found *N*_MG_ to be smaller for the all-wild set than each of the three control sets mentioned in the previous section (blue symbols in **Fig. 3B**). Because each control set is composed of five strains that have exactly the same phylogenetic positions as the five wild strains in the genome tree but share lower environmental similarities than the five wild strains do, the observation of a smaller *N*_MG_ for the all-wild set than each of the control set indicates that having a common environment does not increase *N*_MG_, supporting the neutral hypothesis of expression evolution. Note that the above finding cannot be caused by a potential lack of information in the data, because if we choose six strains from the nine used, the number of genes supporting the monophyly of the six *S. cerevisiae* strains (red dot in **Fig. 3C**) is the highest among all 84 possible six-strain sets (gray dots in **Fig. 3C**). Apparently, our expression dataset contains rich information, but the information points to a neutral rather than adaptive explanation of gene expression variation.

For comparison, we repeated the above analyses using the yeast genome and morphological data, respectively. When using individual gene sequences to reconstruct the trees of the nine strains, we found *N*_MG_ = 33 genes to support the monophyly of the all-wild set (red dot in **Fig. 3D**). Twenty of the 126 possible five-strain sets and 14 of the 15 possible five-strain sets that are composed of the three non-*S. cerevisiae* strains and any two *S. cerevisiae* strains (green and blue symbols in **Fig. 3D**) have *N*_MG_ ≥ 33. *N*_MG_ for the all-wild set is larger than that for one of the three control sets. As expected, a monophyly of the six *S. cerevisiae* strains (red dot in **Fig. 3E**) is supported by more genes than any other six-strain set (grey dots in **Fig. 3E**). Thus, results from the gene sequence data are overall similar to those from the gene expression data. If there is any difference, the expression data appear to support the monophyly of the all-wild set relative to other sets even less often than the sequence data.

On the contrary, for the morphological data, 15 of the 219 traits support the monophyly of the all-wild set (red dot in **Fig. 3F**). This fraction (15/219 = 6.8%) is an order of magnitude greater than the corresponding fractions for genome (33/4325 = 0.76%) and gene expression (22/4325 = 0.51%) data (*P* = 3×10^−9^ and 3×10^−11^, respectively, G-test of independence). Only two of the 15 five-strain sets containing all non-*S. cerevisiae* strains and any two *S. cerevisiae* strains (green and blue symbols **in Fig. 3F**) and 8 of all 126 five-strain sets are supported by ≥ 15 traits. Furthermore, the number of traits supporting the all-wild set exceeds that supporting each of the three control sets (blue symbols in **Fig. 3F**). By contrast, the monophyly of all six *S. cerevisiae* strains, supported by only 27 morphological traits, is no longer the best supported among all six-strain sets.

Altogether, these phylogenetic analyses of individual gene sequences, expression levels, and morphological traits strongly suggest that the observed yeast expression variations within and between species are largely neutral.

### Expression variation among wild strains exceeds that between wild and non-wild strains

If gene expression variations among the yeast strains are mainly due to environmental adaptations, the expression level differences among the five wild strains should be relatively small, compared with those between wild and non-wild strains. On the contrary, if expression variations are mainly neutral, expression differences among strains should increase with their genomic distances. Consequently, under the neutral hypothesis and given the genome tree (**Fig. 1A**), expression differences among wild strains are not expected to be smaller, compared with those between wild and non-wild strains. In other words, the neutral and adaptive hypotheses may also be tested by measuring expression differences of individual genes among various strains without making a tree. We used two measures of gene expression differences. First, for each gene, we calculated the mean *p*air-wise *d*ifference in expression level among all *w*ild strains (PD_ww_), as well as that between all pairs of wild and non-wild strains (PD_wo_). The smaller the ratio between PD_ww_ and PD_wo_, the stronger the evidence for adaptation. Second, for each gene, we also compared the variance in expression level among all wild strains (V_w_) and that among all strains (V_t_). Again, the smaller the ratio between V_w_ and V_t_, the stronger the evidence for adaptation. We plotted the frequency distributions of these two ratios using the actual wild and non-wild strains (**Figs. 4A** and **B**). As three controls, we plotted the frequency distributions when we swap one or both wild *S. cerevisiae* strains with their respective non-wild sister strains. But, neither PD_ww_/PD_wo_ nor V_w_/V_t_ is smaller for the actual wild strains when compared with the three controls (*P* > 0.5, one-tail Mann–Whitney *U* test; **Figs. 4A** and **B**), consistent with the neutral hypothesis.

**Figure 4.**
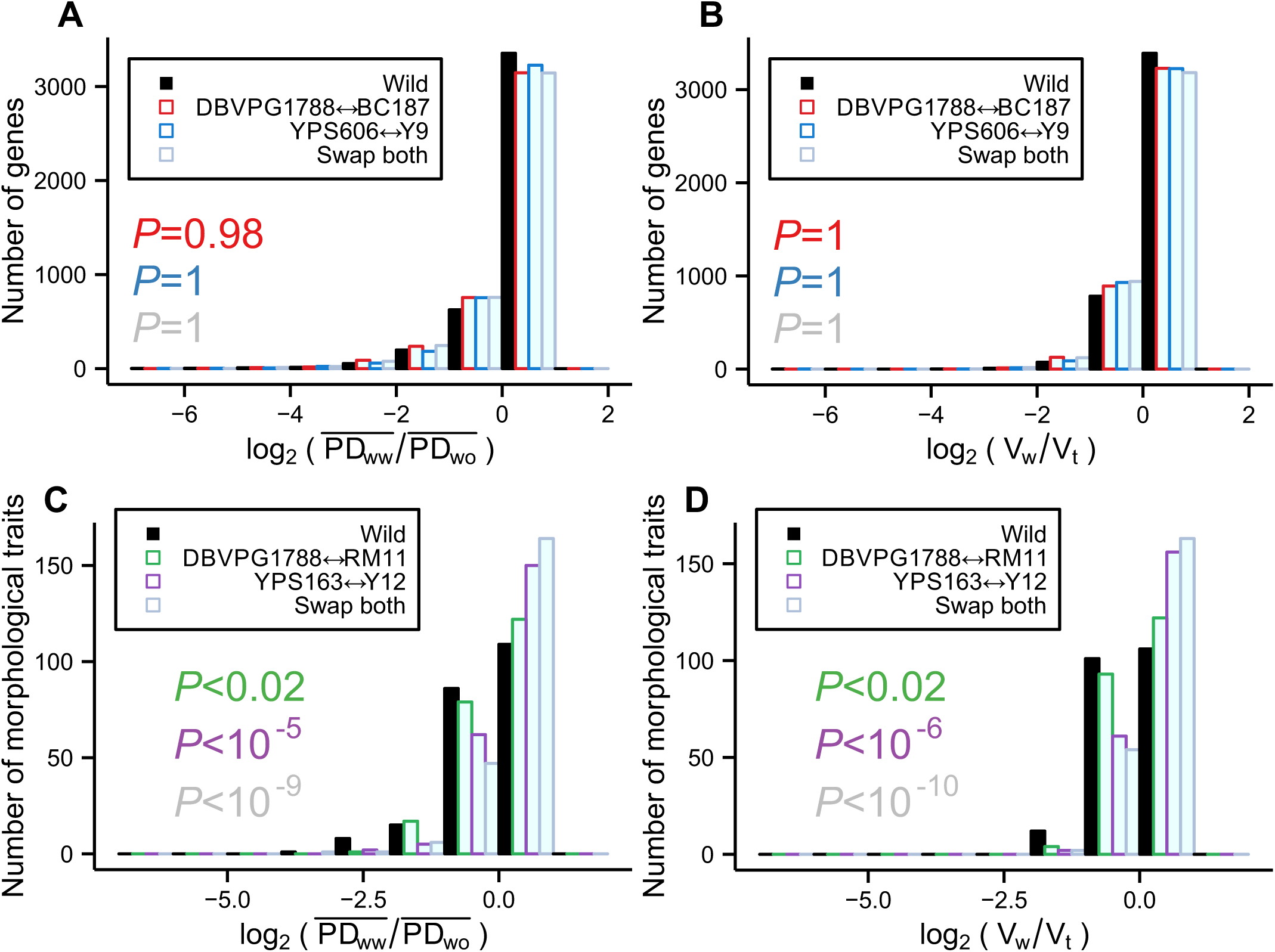
Gene expression variances among wild strains and among all strains. (A) Frequency distribution of the logarithm of the ratio between the mean expression difference between wild strains and that between wild and non-wild strains (black). As controls, the same quantity is plotted when one or both of the wild *S. cerevisiae* strains are swapped with their sister non-wild strains in the calculation. The *P-*value from Mann–Whitney *U* test, measuring the probability that the median value of the observed distribution (black) is equal to or greater than that of a control distribution, is indicated with the same color as the control distribution. (B) Frequency distribution of the logarithm of the ratio between the variance in expression level among the wild strains and that among all strains (black). As controls, the same quantity is plotted when one or both of the wild *S. cerevisiae* strains are swapped with their sister non-wild strains in the calculation. The *P-*value from Mann–Whitney *U* test, measuring the probability that the median value of the observed distribution is equal to or greater than that of a control distribution, is indicated with the same color as the control distribution. (C) Same as A except that morphological data instead of gene expression data are used. (D) Same as B except that morphological data instead of gene expression data are used.

For comparison, we performed the same analysis with the morphological data. Interestingly, both PD_ww_/PD_wo_ and V_w_/V_t_ are significantly skewed towards lower values when compared with the three controls (**Figs. 4C and D**), consistent with the adaptive evolution hypothesis of morphological traits (19).

## DISCUSSION

In this study, we measured genome-wide gene expression levels in nine yeast strains belonging to three closely related species and isolated from five different ecological environments. We repeatedly found that the intra- and inter-specific variations in gene expression levels can be explained by the neutral accumulation of random mutations but are inconsistent with the environmental adaptation hypothesis. Our study has several caveats that are worth considering.

First, the quality of the gene expression data is critical to our conclusion. The sequencing depth in our RNA-seq experiment is high, with ∼70.4 million mapped reads per sample. For the gene with the median expression level in our data, there are on average 93 reads covering each nucleotide. Further, biological replicates show highly similar expressions (**Fig. S1**) and form a monophyly in transcriptome trees (**Fig. S3**). That the transcriptome tree becomes topologically even closer to the genome tree after the removal of 25% to 50% lowly expressed genes suggests that our conclusion of neutral evolution drawn from analyzing the whole transcriptome data is conservative.

Second, the laboratory condition under which gene expressions are quantified is important to the test of the neutral and adaptive hypotheses. The oak exudate medium used mimics the natural habitat of wild yeast strains; our previous study showed that, compared with some non-wild strains, a wild strain grows faster in this medium than in the other commonly used media tested (23). Thus, using this medium should help reveal environmental adaptations of gene expressions in the wild strains if such adaptations exist. The absence of adaptive signals even in this medium implies the unlikelihood of detecting adaptations in other conditions that are less similar to the natural habitat of the wild strains. Nonetheless, the synthetic oak exudate medium is not identical to the natural habitat of the wild yeast strains, which could have obscured the potential adaptive signals in gene expression variations. However, natural environments fluctuate and gene expression evolution includes the evolution of expression responses to environmental fluctuations, of which the oak exudate medium may be considered one. If gene expression variations are primarily adaptive, one cannot explain why the among-strain variation in the expression response to the oak exudate medium is structured like the genome tree of the strains. By contrast, this pattern of variation is expected if it is due to the random accumulation of neutral mutations. Thus, although the medium used in our experiment is not identical to the natural habitat of the wild yeast strains, our findings regarding the evolutionary mechanism of expression variations are informative. Of course, this conclusion should be further verified under other relevant conditions in the future.

Third, although the ecological environments of the five wild strains are overall similar, these environments may still differ in temperature, humidity, day length, etc. Consequently, one could argue that the adaptive hypothesis does not necessarily predict similar gene expressions among the wild strains. While this argument may be valid, the adaptive hypothesis cannot explain the significant topological similarity between the transcriptome tree and the genome tree, because the differences among the environments of the nine strains are certainly not represented by the genome tree. The similarity between the transcriptome and genome trees, in contrast to the dissimilarity between the morphology and genome trees, strongly supports the neutral explanation of the expression variations among the nine yeast strains, particularly in the light of the recent finding of adaptive variations of the morphological traits examined (19).

Fourth, we assumed that environmental adaptation means that there is a single optimal expression level or a continuous range of equally optimal expression levels for a given gene in an environment. This assumption, however, may not be correct for all genes. Strains from the same ecological environment but with different genetic backgrounds could have different optimal gene expression levels, due to different genetic interactions that have accumulated since the strains diverged from one another. For example, let X and Y be two genes with virtually identical functions and let the optimal total expression level of X and Y in the natural yeast habitat be anywhere between 10 and 12 units. Under this scenario, the optimal expression level of X will vary among wild strains if these strains have different expression levels of Y. If this or other scenarios of genetic interactions apply to most genes in the genome, our analysis would be incapable of testing the adaptive hypothesis, because what is potentially selectively optimized is not the expression levels of individual genes but unknown mathematical functions of the expression levels of multiple genes. However, for the following reason, such genetic interactions cannot be widespread. We previously found that deleting certain genes in a lab strain of yeast can increase the yeast fitness under a given environment, which led us to predict that these genes should have been down-regulated in strains well adapted to that environment (23). Interestingly, this prediction is usually correct (23), suggesting that genes with fitness effects upon deletion are rarely subject to the type of genetic interaction aforementioned, because otherwise the prediction could not have been so good. This consideration suggests that the necessary assumption required for our rejection of the adaptive hypothesis is likely satisfied for most genes.

While our study suggests the paucity of adaptive expression variations among the yeast strains studied, it does not exclude the possibility of a small fraction of genes whose expression variations are largely caused by adaptations. In fact, adaptive expressional differences among *S. cerevisiae* strains have been suggested for some genes (23-25). It should be pointed out, however, that identifying an adaptive signal in the expression difference of a gene between two strains does not necessarily mean that their expression difference is entirely or even largely explained by adaptation. The following hypothetical example illustrates this point. The optimal expression level for a gene in strain A is anywhere between 5 and 10 units, while its optimal expression level in strain B is anywhere between 25 and 100 units. If we observe that the expression level of the gene is 7 units in strain A and 82 units in strain B, the fraction of their expression difference that is potentially adaptive is only (25−10)/(82−7) = 20%. Adaptive signals can be detected in some tests even when the fraction of expression variation explainable by adaptation is low (23), whereas some seemingly adaptive patterns, such as the similarity in the expression levels of the 22 genes among the wild strains in Fig. 3A, are better explained by neutral evolution upon a closer examination. If only a small fraction of expression differences between two populations are adaptive, the adaptive expression differences likely arise early in their environmental adaptations, because the fitness advantages of successive fixations of mutations in an adaptation process are expected to decline exponentially (26). It is thus not surprising that three parallel populations of yeast under continuous aerobic growth in glucose-limited chemostats for 250-500 generations showed parallel expression changes for some genes (27), although it is unclear whether the number of genes with parallel expression changes significantly exceeds the random expectation. Notably, even during early stages of adaptations, the amount of adaptive expression changes may still be limited, compared with neutral changes. For example, in a large-scale experimental evolution study that subjected eight populations of yeast to each of three conditions (glucose limitation, sulfate limitation, and phosphate limitation) for 100-400 generations of mitotic growth, populations selected under the same conditions do not form monophyletic clades in the transcriptome tree (28). Regardless, these experimental evolution studies were not designed to distinguish between the neutral and adaptive hypotheses and hence do not provide critical evidence for or against each hypothesis.

Recently, Karen et al. measured the fitness effects on a laboratory yeast strain by individually altering the expression levels of ∼100 genes (29). Due to the limited sensitivity of the laboratory fitness quantification, a measured fitness effect <1% cannot be distinguished from no effect. We thus estimated the "neutral" expression range, for which the measured fitness effect is no greater than 1%. For 69% of the 78 genes that can be analyzed (29), the neutral expression range is wider than the actual expression range observed in the nine strains studied here, explaining why the observed expression variations within and between species appear neutral for most genes. The above comparison is of course approximate, because the media used in the laboratory fitness measurement differ from that used in our study, the genetic background stays unchanged in the fitness measurement but varies among our nine strains, and the sensitivity of the fitness assay is lower than that of natural selection. Nevertheless, the comparison shows that the direct fitness measures are not inconsistent with our conclusion.

Except for one early microarray study that suggested no purifying selection (11), there is ample evidence for and general agreement on the action of purifying selection in gene expression evolution (30-33). That early suggestion was based on a lack of significant difference in the rate of expression evolution between intact genes and expressed pseudogenes (11), which could have been due to low statistical power caused by the inclusion of too few expressed pseudogenes and/or the action of purifying selection on expressed pseudogenes (34-36). While our study is not intended to detect purifying selection in gene expression evolution, our data are consistent with the action of purifying selection. For instance, for each gene, we measured the coefficient of variation in expression level among all nine strains (CV_t_) and that among the five wild strains (CV_w_), and found both to be negatively correlated with the importance of the gene measured by the fitness reduction caused by deleting the gene in the oak exudate medium (23) (for CV_t_: ρ = −0.095, *P* = 0.0001; for CV_w_: ρ = −0.081, *P* = 0.0009). This pattern is explainable by stronger purifying selection acting on the expression levels of more important genes.

After the removal of deleterious expression variations by purifying selection, the remaining expression variations that are observed can be neutral or adaptive. We found that they are largely neutral rather than adaptive. Given the relatively high expression variations observed (mean CV_t_ = 0.39 and mean CV_w_ = 0.36), this conclusion seems to be at odds with the view that gene expression levels are tightly regulated and consequently should show little neutral variation. For instance, an elegant experiment on the *Escherichia coli* lactose operon suggests that protein expression levels are finely tuned according to the cost and benefit of gene expression and protein production, which vary depending on the environment (37). Similarly, by considering the energy cost of protein synthesis, Wagner estimated that natural selection would prohibit a >2% increase in protein concentration above the optimal level for any gene that is more highly expressed than the median gene expression level in yeast (38). One possibility that could potentially resolve the apparent conflict between these findings and our results is the existence of posttranscriptional regulations that minimize the downstream consequences of variations in mRNA concentrations. Indeed, several studies have shown that protein concentrations are generally more conserved evolutionarily than mRNA concentrations (39, 40) and that mRNA concentration differences between species are often offset by differences in translation (41-43).The much smaller energy cost of mRNA synthesis than that of protein synthesis (38) also permits a larger range of neutral variation in mRNA concentration. These considerations lead us to hypothesize that the adaptive fraction of intra- and inter-specific variations in protein concentration is greater than the adaptive fraction of gene expression variations. With the rapid progress of quantitative proteomics, this hypothesis may be tested in the near future.

In terms of how directly a trait impacts the organism-level phenotype, the four types of traits discussed in this work can be ranked as organismal morphology, protein concentration, mRNA concentration, and genome sequence. Because natural selection acts at the organism level, it seems plausible that the more directly that a trait affects the organism-level phenotype, the higher the probability that adaptation contributes to its natural variation. This hypothesis is supported by the present study and can be further tested when comparative proteomic data aforementioned become available.

The role of stochasticity in genotypic evolution is well recognized, while that in phenotypic evolution is less appreciated and agreed upon. Our finding that natural variations in gene expression level, a molecular phenotype, is generally shaped by stochastic genetic drift rather than deterministic adaptation expands the role of stochasticity in evolution. It is likely that the role of stochasticity in evolution, compared with that of adaptation, is generally reduced as one moves from traits that impact the organismic phenotype less directly to those that impact more directly.

It has been heatedly debated whether phenotypic adaptations seen at the organism level are mainly caused by protein sequence/function changes or gene expression changes, especially those brought about by alterations of *cis*-regulatory sequences (10, 44). We previously provided evidence supporting the hypothesis that evolution of morphological traits is more often caused by gene expression changes while that of physiological traits is more often owing to protein function changes (45). Regardless, our finding that gene expression variations are largely neutral should reduce our expectation that an organismic adaptation is caused by expression changes (see Materials and Methods).

Given the huge effective population size of yeast, our finding that yeast expression variations are largely neutral suggests that the same would apply to species with smaller effective population sizes, which include almost all multicellular organisms. Some early microarray-based gene expression studies from invertebrates (46) and vertebrates (47) reported at most a tiny fraction of adaptive expression variations and are thus consistent with our prediction, but a stronger test of neutral expression variation is warranted. Note that the observation (31, 48) that the transcriptomes of multiple organs from several mammalian species are clustered by organ rather than species (e.g., human liver transcriptome is closer to mouse liver transcriptome than to human heart transcriptome) does not distinguish between the neutral and adaptive hypotheses, because this clustering is predicted by both hypotheses due to the fact that different organs originated prior to the emergence of mammals and that they have distinct functions. It will be of great interest and importance to test the prediction that intra- and inter-specific expression variations in multicellulars are mostly neutral.

## MATERIALS AND METHODS

### Yeast genome sequences and phylogenetic analysis

The genome sequences of all *S. cerevisae* strains used here (Table S1) were obtained from a recently completed yeast population genomic study that sequenced 85 *S. cerevisiae* strains from a diverse array of ecological and geographic origins (21). *S. cerevisiae* genomic annotations were downloaded from SGD (22). The two *S. paradoxus* and one *S. mikatae* genome sequences and their annotations were previously published (49). We first identified *r*eciprocal *b*est *h*its (RBH) between *S. cerevisiae* and each of the other two species in a specie-wise tBLASTx search (50) among all annotated genes, using an *E*-value cutoff of 10^−4^. To avoid the complication of gene expression changes after gene duplication, we should exclude paralogs generated after the separation of the three species and include only one-to-one orthologs among the species. To this end, we removed from the above RBH gene list any gene that is the best hit of a gene from either of the other two yeasts but not on the list, resulting in a set of 4625 one-to-one orthologous genes. We further removed those genes that contain undetermined nucleotides in coding regions due to incomplete genomic sequencing. Our final list had 4325 genes.

We aligned the coding sequences of each of the 4325 genes from the nine yeast strains by MACSE (51) and removed alignment gaps. The aligned sequences of each gene were then used by PHYLIP (52) to estimate F84 pairwise nucleotide distances and reconstruct a neighborjoining (NJ) tree (53) of the nine strains. To reconstruct the genome tree, we first concatenated the coding sequence alignments of all genes and then estimated F84 distances and built an NJ tree using PHYLIP. Statistical support for each interior branch of the genome tree was assessed by bootstrapping the 4325 genes 1000 times.

### RNA sequencing and transcriptome analysis

Each of the nine yeast strains was streaked to form single colonies from frozen glycerol stocks held at −80°C onto YPD plates (1% yeast extract, 2% peptone, 2% glucose, 2% agar). After 48 hours of growth at 30°C, a single colony was picked and inoculated into 5 ml of the synthetic oak exudate medium (1% sucrose, 0.5% fructose, 0.5% glucose, 0.1% yeast extract, and 0.15% peptone) (14). Strains were grown for 24 hours at 30°C before dilution into fresh synthetic oak exudate medium to an OD660 of 0.1. Cultures were grown at 30°C until OD660 = 0.5 (mid-log phase), at which point cells were harvested by centrifugation. RNA-seq libraries were prepared following a previous study (54). Briefly, total RNA was extracted from each population using RiboPure-Yeast Kit (Ambion) and treated with DNase I to remove any contaminant DNA. Extraction of mRNA was carried out using MicroPoly(A)Purist Kit (Ambion) and 200 ng of the resulting mRNA sample was fragmented (Fragmentation Buffer, NEB) before ethanol precipitation. First strand cDNA synthesis was performed using random hexamer priming (Superscript II, Invitrogen), followed by second strand cDNA synthesis (Invitrogen) as recommended by the manufacturer. End repair, A-tailing, and ligation of the Illumina adapters necessary for sequencing were then carried out using the NEBnext mRNA sample preparation kit (NEB). Libraries were then size-selected by agarose gel electrophoresis followed by gel extraction such that libraries consisted of fragments containing inserts of approximately 250 bp in length. Polymerase-chain-reaction amplification was performed for 15 cycles using NEBnext mRNA sample preparation kit, before single-end sequencing on the Illumina GAII platform was performed at the University of Michigan Sequencing Core. Sequencing statistics are listed in Table S1.

All raw read sequences generated by RNA-seq were first processed by cutadapt (55) to trim any remaining adaptor sequences. The trimmed reads were then aligned to the genome of the corresponding strains by tophat (56) under the default parameter set except that a maximal intron size of 10 kb was allowed because the largest annotated intron in *S. cerevisiae* is 9349 bp (22). Alignment results were fed to cufflinks (56) for quantification of known transcripts in *S. cerevisiae* (22), *S. paradoxus* (49), and *S. mikatae* (49). Unless otherwise noted, all gene expression levels used in our analyses are in the unit of RPKM (*r*eads *p*er *k*ilobases per *m*illion reads). All RNA-seq reads as well as estimated gene expression levels have been deposited in NCBI with a GEO ID of GSE81320.

The expression levels of the 4325 genes used for constructing the genome tree were analyzed. These genes each have at least one RNA-seq read in at least one of the nine strains. Let *X*_*ij*_ be the expression level in RPKM of gene *i* in strain *j* and *X*_*i*_ be the mean expression level of gene *i* in the nine strains. The Euclidian distance in expression level of gene *i* between strains *j* and *k* is defined by 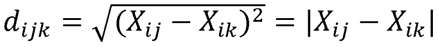 We then used this distance measure to build the NJ tree of the nine strains for gene *i*.

To analyze the transcriptome data as a whole, for each gene *i*, we converted the raw expression levels of the nine strains to standardized expression levels by *Y*_*ij*_= (*X*_*ij*_ − *X*_*i*_)/*S*_*i*_, where *S*_*i*_ is the standard deviation of the expression level of gene *i* among the nine strains. We calculated the transcriptomic Euclidian distance between strains *j* and *k* using the standardized expression levels of all 4325 genes by 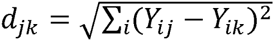.We then built the NJ tree using these distances. The confidence of the transcriptome tree was assessed by bootstrapping the 4325 genes 10,000 times. We similarly built an expression tree for each GO using the standardized Euclidian distances calculated from the standardized expression levels of the genes belonging to the GO.

### Yeast morphological data and analysis

The data of 219 morphological traits from nine strains were obtained from two studies (18, 19). The original data contained 220 traits (19), but one of them (trait ID A103_C) was not used because the data from strain YJM145 were missing. The phylogenetic analysis using these traits was conducted in exactly the same way as the analysis using the gene expression data. The NJ trees for the expression and morphological data were built using the APE package (57).

### Generation of random trees

We generated random trees (in terms of topology) of the nine strains by repeatedly clustering two randomly chosen strains at a time until all nine strains are clustered; after two strains are clustered, they together are considered as a strain in the next round of clustering.

### Topological distance between two trees

Given an unrooted tree structure, each (internal or external) branch connects two sets of tips. In other words, each branch represents a bipartition of the tips. The topological distance between two unrooted trees of the same set of tips is twice the number of internal branches defining different bipartitions of the tips between the two trees (58).

### Principal component analysis (PCA)

To perform PCA of the genome sequences of the nine strains, we used the concatenated multiple sequence alignment of the coding sequences of all 4325 genes. Each site of each sequence in the alignment was converted to a four dimensional vector, whose four components are assigned 1 (or 0) based on the appearance (or not) of A, C, G, and T at this site, respectively. In other words, a sequence with *L* nucleotides was converted to a vector of length 4*L*. The alignment of the nine strains was converted to nine vectors with "aligned" components. For each gene or morphological trait, the expression levels or morphological trait values from the nine strains were first scaled to the standard normal distribution before PCA. PCA was conducted using the "prcomp" function in the "stats" package in R (59).

### Posterior probabilities of protein function and gene expression changes

The posterior probability that an organismic phenotypic adaptation is caused by an expression change, *P*(E|A), relative to the posterior probability that it is caused by a protein function change, *P*(F|A), can be calculated by [*P*(A|E)/*P*(A|F)][*P*(E)/*P*(F)] according to the Bayes’ theorem, where *P*(E) and *P*(F) are the prior probabilities of expression changes and protein function changes, respectively, and *P*(A|E) and *P*(A|F) are the probabilities of having an organismic phenotypic adaptation conditional on an expression change and a protein function change, respectively. Our result that *P*(A|E) is smaller than previously thought reduces the expectation that an organismic adaptation is caused by a gene expression change.

## ACKNOWLEDGEMENTS

This work was supported by the research grant R01GM103232 from the U.S. National Institutes of Health to J.Z.

**Figure S1.** High correlation (*R*) between the gene expression levels of strain YPS606 measured in two biological replicates. Each dot is a gene. RPKM, reads per kilobases per million reads.

**Figure S2.** Transcriptome trees showing the two biological replicates of YPS606. Panels (A)-(C) are equivalent to Fig. 1B-D, respectively, except that the data from the two biological replicates of YPS606 are not merged.

**Figure S3**. Principal component analysis of gene expression levels of the nine strains. (A-B) Same as Fig. 2B, except that the (A) 75% or (B) 50% most highly expressed genes instead of all genes are used.

(C) Frequency distribution (bars) of the mean distance between two *S. cerevisiae* strains (*d*_*Sc-Sc*_) divided by the mean distance between a *S. cerevisiae* strain and a *S. paradoxus* strain (*d*_*Sc-Sp*_) in the transcriptome PCA plot made with 1000 random sets of 219 genes. The arrow shows the corresponding ratio in the morphology PCA plot with 219 traits.

## REFERENCES

1. Darwin C (1859) On the Origin of Species by Means of Natural Selection (J. Murray, London,).

2. Futuyma DJ (2013) Evolution (Sinauer Associates) 3rd Ed.

3. Mayr E (2001) What Evolution is (Basic Books, New York).

4. Endler JA (1986) Natural Selection in the Wild (Princeton University Press, Princeton, N.J.).

5. Nei M (2013) Mutation-Driven Evolution (Oxford University Press, Oxford).

6. Nei M (1987) Molecular Evolutionary Genetics (Columbia University Press, New York).

7. Lynch M (2007) The Origins of Genome Architecture (Sinauer, Sunderland, Mass).

8. Kimura M (1968) Evolutionary rate at the molecular level. Nature 217(5129):624–626.

9. Carroll SB (2008) Evo-devo and an expanding evolutionary synthesis: a genetic theory of morphological evolution. Cell 134(1):25–36.

10. Stern DL & Orgogozo V (2008) The loci of evolution: how predictable is genetic evolution? Evolution 62(9):2155–2177.

11. Khaitovich P, et al. (2004) A neutral model of transcriptome evolution. PLoS Biol 2(5)E132.

12. Gilad Y, Rifkin SA, Bertone P, Gerstein M,&White KP (2005) Multi-species microarrays reveal the effect of sequence divergence on gene expression profiles. Genome Res 15(5):674–680.

13. Sniegowski PD, Dombrowski PG, & Fingerman E (2002) Saccharomyces cerevisiae and Saccharomyces paradoxus coexist in a natural woodland site in North America and display different levels of reproductive isolation from European conspecifics. FEMS Yeast Res 1(4):299–306.

14. Murphy HA, Kuehne HA, Francis CA,&Sniegowski PD (2006) Mate choice assays and mating propensity differences in natural yeast populations. Biol Lett 2(4):553–556.

15. Johnson LJ, et al. (2004) Population genetics of the wild yeast Saccharomyces paradoxus. Genetics 166(1):43–52.

16. Wang QM, Liu WQ, Liti G, Wang SA,&Bai FY (2012) Surprisingly diverged populations of Saccharomyces cerevisiae in natural environments remote from human activity. Mol Ecol 21(22):5404–5417.

17. Warringer J, et al. (2011) Trait variation in yeast is defined by population history. PLoS Genet 7(6) e1002111.

18. Yvert G, et al. (2013) Single-cell phenomics reveals intra-species variation of phenotypic noise in yeasts. BMC Syst Biol 7:54.

19. Ho W-C, Ohya Y,&J. Zhang (2016) Testing the neutral hypothesis of phenotypic evolution. BioRxiv doi: http://dx.doi.org/10.1101/089987.

20. Liti G, et al. (2009) Population genomics of domestic and wild yeasts. Nature 458(7236):337–341.

21. Maclean CJ, et al. (2016) Deciphering the genic basis of environmental adaptations by simultaneous forward and reverse genetics in Saccharomyces cerevisiae. BioRxiv doi: http://dx.doi.org/10.1101/087510.

22. Cherry JM, et al. (2012) Saccharomyces Genome Database: the genomics resource of budding yeast. Nucleic Acids Res 40(Database issue):D700–705.

23. Qian W, Ma D, Xiao C, Wang Z,&Zhang J (2012) The genomic landscape and evolutionary resolution of antagonistic pleiotropy in yeast. Cell Rep 2(5):1399–1410.

24. Fraser HB, Moses AM,&Schadt EE (2010) Evidence for widespread adaptive evolution of gene expression in budding yeast. Proc Natl Acad Sci U S A 107(7):2977–2982.

25. Bullard JH, Mostovoy Y, Dudoit S,&Brem RB (2010) Polygenic and directional regulatory evolution across pathways in Saccharomyces. Proc Natl Acad Sci U S A 107(11):5058–5063.

26. Orr HA (1998) The population genetics of adaptation: the distribution of factors fixed during adaptive evolution. Evolution 52:935–949.

27. Ferea TL, Botstein D, Brown PO,&Rosenzweig RF (1999) Systematic changes in gene expression patterns following adaptive evolution in yeast. Proc Natl Acad Sci U S A 96(17):9721–9726.

28. Gresham D, et al. (2008) The repertoire and dynamics of evolutionary adaptations to controlled nutrient-limited environments in yeast. PLoS Genet 4(12) e1000303.

29. Keren L, et al. (2016) Massively parallel interrogation of the effects of gene expression levels on fitness. Cell 166(5):1282–1294.

30. Denver DR, et al. (2005) The transcriptional consequences of mutation and natural selection in Caenorhabditis elegans. Nat Genet 37(5):544–548.

31. Liao BY & Zhang J (2006) Evolutionary conservation of expression profiles between human and mouse orthologous genes. Mol Biol Evol 23(3):530–540.

32. Rifkin SA, Houle D, Kim J,&White KP (2005) A mutation accumulation assay reveals a broad capacity for rapid evolution of gene expression. Nature 438(7065):220–223.

33. Jordan IK, Marino-Ramirez L,&Koonin EV (2005) Evolutionary significance of gene expression divergence. Gene 345(1):119–126.

34. Podlaha O & Zhang J (2010) Pseudogenes and their evolution. Encyclopedia of Life Sciences, (John Wiley&Sons, Chichester, UK), pp 1–8.

35. Xu J & Zhang J (2016) Are human translated pseudogenes functional? Mol Biol Evol 33(3):755–760.

36. Khachane AN & Harrison PM (2009) Assessing the genomic evidence for conserved transcribed pseudogenes under selection. BMC Genomics 10:435.

37. Dekel E & Alon U (2005) Optimality and evolutionary tuning of the expression level of a protein. Nature 436(7050):588–592.

38. Wagner A (2005) Energy constraints on the evolution of gene expression. Mol Biol Evol 22(6):1365–1374.

39. Laurent JM, et al. (2010) Protein abundances are more conserved than mRNA abundances across diverse taxa. Proteomics 10(23):4209–4212.

40. Schrimpf SP, et al. (2009) Comparative functional analysis of the Caenorhabditis elegans and Drosophila melanogaster proteomes. PLoS Biol 7(3) e48.

41. Artieri CG & Fraser HB (2014) Evolution at two levels of gene expression in yeast. Genome Res 24(3):411–421.

42. McManus CJ, May GE, Spealman P,&Shteyman A (2014) Ribosome profiling reveals post-transcriptional buffering of divergent gene expression in yeast. Genome Res 24(3):422–430.

43. Khan Z, et al. (2013) Primate transcript and protein expression levels evolve under compensatory selection pressures. Science 342(6162):1100–1104.

44. Hoekstra HE & Coyne JA (2007) The locus of evolution: evo devo and the genetics of adaptation. Evolution 61(5):995–1016.

45. Liao BY, Weng MP, & Zhang J (2010) Contrasting genetic paths to morphological and physiological evolution. Proc Natl Acad Sci U S A 107(16):7353–7358.

46. Rifkin SA, Kim J, & White KP (2003) Evolution of gene expression in the Drosophila melanogaster subgroup. Nat Genet 33(2):138–144.

47. Oleksiak MF, Churchill GA, & Crawford DL (2002) Variation in gene expression within and among natural populations. Nat Genet 32(2):261–266.

48. Brawand D, et al. (2011) The evolution of gene expression levels in mammalian organs. Nature 478(7369):343–348.

49. Scannell DR, et al. (2011) The Awesome Power of Yeast Evolutionary Genetics: New Genome Sequences and Strain Resources for the Saccharomyces sensu stricto Genus. G3 (Bethesda) 1(1):11–25.

50. Camacho C, et al. (2009) BLAST+: architecture and applications. BMC Bioinformatics 10:421.

51. Ranwez V, Harispe S, Delsuc F, & Douzery EJ (2011) MACSE: Multiple Alignment of Coding SEquences accounting for frameshifts and stop codons. PLoS One 6(9) e22594.

52. Felsenstein J (1989) PHYLIP - Phylogeny Inference Package (Version 3.2). Cladistics 5:164–166.

53. Saitou N & Nei M (1987) The neighbor-joining method: a new method for reconstructing phylogenetic trees. Mol Biol Evol 4(4):406–425.

54. Nagalakshmi U, et al. (2008) The transcriptional landscape of the yeast genome defined by RNA sequencing. Science 320(5881):1344–1349.

55. Martin M (2011) Cutadapt removes adapter sequences from high-throughput sequencing reads. EMBnet.journal 17:10–12.

56. Pollier J, Rombauts S, & Goossens A (2013) Analysis of RNA-Seq data with TopHat and Cufflinks for genome-wide expression analysis of jasmonate-treated plants and plant cultures. Methods Mol Biol 1011:305–315.

57. Paradis E, Claude J, & Strimmer K (2004) APE: Analyses of Phylogenetics and Evolution in R language. Bioinformatics 20(2):289–290.

58. Penny D & Hendy MD (1985) The Use of Tree Comparison Metrics. Systematic Biology 34:75–82.

59. R. Core Team (2013) R: A Language and Environment for Statistical Computing (R Foundation for Statistical Computing, Vienna, Austria).

